# Evaluation of Gene Expression and Phenotypic Profiling Data as Quantitative Descriptors for Predicting Drug Targets and Mechanisms of Action

**DOI:** 10.1101/580654

**Authors:** Maris Lapins, Ola Spjuth

**Affiliations:** Department of Pharmaceutical Biosciences, Uppsala University, Uppsala, Sweden

## Abstract

Profiling drug leads by means of in silico and in vitro assays as well as omics is widely used in drug discovery for safety and efficacy predictions. In this study, we evaluate the performance of machine learning models trained on data from gene expression and phenotypic profiling assays, with models trained on chemical structure descriptors, for prediction of various drug mechanisms of action and target proteins. Models for several hundred mechanisms of actions and targets were trained using data on 1484 compounds characterized in both gene expression using L1000 profiles, and phenotypic profiling with cell painting assay. The results indicate that the accuracy of the three profiling technologies varies for different endpoints, and indicate a clear potential synergistic effect if these methods are combined. We also study the effect of predictive accuracy of data from different cell lines for L1000 profiles, showing that the choice of cell line has a non-negligible effect on the predictive accuracy. The results strengthen the idea of integrated approaches for predicting drug targets and mechanisms of action in preclinical drug discovery.

## Introduction

Over the past decade, methods have been developed to systematically determine cellular effects of chemical compounds with the aim to improve fields such as drug screening and safety profiling.^1–3^ Important objectives include to predict off-target effects and adverse drug reactions but also to offer insights into compound’s mode of action and the establishment of adverse outcome pathways.

Pharmaceutical profiling using ligand binding or enzyme assays is the most widely used in vitro methodology, and it is widely implemented in drug discovery safety platforms. Profiling using gene expression is relatively recent, and pioneering work includes Connectivity Map^4^ that has been widely used and built upon.^5,6^

L1000 is a high throughput and low cost gene expression profiling method, based on representation of transcriptome by 978 “landmark genes”. Recently, datasets with L1000 profiles were made available in Broad LINCS L1000 Connectivity Map project, including profiles for a total of 20K small molecule compounds, of which over 2K compounds were studied systematically in nine human cancer cell lines.^6,7^

Multiparametric high-content imaging has also proven to be a highly useful and successful technique for understanding biological activity in response to chemical and genetic perturbations. The Broad Bioimage Benchmark Collection (BBBC) is an important publicly available collection of microscopy images. Some of the largest image sets obtained by Cell Painting assay comprise osteosarcoma cells treated by 1.6K known bioactive compounds^8^ and by 30K compounds, most of which being derived from diversity-oriented synthesis.^9^

It is hypothesized that chemical compounds with a similar mechanism of action (MoA), which act upon the same signaling pathways, will produce comparable phenotypes, and that analysis of phenotyping profiling data can predict compound mechanism of action.^10^ Successful prediction examples include a study by Ljosa et al.^11^ where 37 compounds are classified to 12 MoA’s with 94% prediction accuracy, and a study by Warchal et al. where 24 compounds are classified to eight MoA’s with over 80% accuracy in several cell lines.^10^ On a large scale, predicting of results of particular biological assays on the basis of phenotyping profiling data have been recently undertaken.^12,13^ In particular, in a study by Simm et al. information extracted from microscopy-based screen for glucocorticoid receptor translocation was able to predict assay-specific biological activity in two ongoing drug discovery projects, leading to a tremendous 60-fold and 250-fold increase of hit rates.

For transcriptomic data, models are reported by Aliper et al.^14^ where several hundred compounds selected from Broad LINCS database are linked to 12 therapeutic use categories in breast cancer (MCF7), prostate cancer (PC3), and lung cancer cells (A549).

The aim of the current study was to compare the performances of features derived from gene expression and phenotypic profiling assays with the performance of chemical structure based descriptors, for prediction of various drug mechanisms of action and target proteins. To this end, models for several hundred mechanism(s) of actions and/or target(s) (MoA/Ts) were created using data for 1484 compounds characterized in both gene expression and phenotypic profiling assays. As L1000 gene expression profiles have been collected systematically in several cell lines, we also aimed to investigate cell-context specificity of transcriptomic data for predicting MoA/Ts.

In phenotyping profiling, each compound is typically tested in quadruplicates or octaplicates on different plates and thus four or eight profiles per compound are obtained. The overall profile thus depends on the way the data are aggregated. In this study we therefore also investigated the effects of data pre-processing on the prediction accuracy.

## Materials and Methods

### Datasets

#### Gene expression profiling (Connectivity Map)

The Connectivity Map (CMap) dataset built using L1000 high-throughput gene-expression assay was downloaded from Gene Expression Omnibus (ascension: GSE92742).^7^ The dataset comprises transcriptional responses (expression of 978 landmark genes) to perturbations of various cells by 19,811 small molecule compounds. 2,429 of the compounds are profiled systematically across nine human cancer cell lines. Most of the profiles are obtained with 10 μM dose and 24 hour treatment time.

#### Phenotypic profiling (Cell Painting)

Dataset of images and morphological profiles of 30,616 small molecule treatments obtained by Cell Painting assay was downloaded from GigaDB, http://gigadb.org/dataset/100351. In this assay, human U2OS (human osteosarcoma) cells are stained for eight major organelles and sub-compartments, using a mixture of six fluorescent dyes.^15^ From five channel microscopy images, 1783 morphological features are generated by CellProfiler software.^16^

### Compound annotation with protein targets and/or mechanism of action

We used Touchstone data base (https://clue.io/touchstone)^6^ and Drug Repurposing Hub (https://clue.io/repurposing)^17^ to associate compounds to their mechanism(s) of actions and/or protein target(s) (MoA/Ts). From annotations to individual targets, we also derived labels for protein kinase groups.

For Cell Painting dataset, we obtained annotations for 1759 compounds, where 262 MoA/Ts were shared by at least five compounds. In CMap dataset, the three cell lines with the highest number of annotated compounds were MCF7 (breast cancer, 2801 annotated compounds, 444 MoA/Ts shared by at least five compounds), PC3 (prostate cancer, 2775 annotated compounds, 435 MoA/Ts), and A549 (lung cancer, 2319 annotated compounds, 380 MoA/Ts). The intersection of Cell Painting dataset and the largest of CMap datasets (MCF7) contained 1484 compounds and 234 MoA/Ts.

### Data pre-processing

In Cell Painting dataset, most of the compounds have been used to treat cells eight times on different plates, thus giving eight sets of morphological features for each compound. In data pre-processing, we first centered and (optionally) normalized the features on plate-to-plate basis, by subtracting the mean value and (optionally) dividing by the standard deviation for the control samples on this plate. Thereafter we calculated the mean or the median values of each feature from the eight sets, and used them as descriptors for the compounds. Some of the 1783 features were invariant in the present dataset, and were removed before the modelling.

### Random Forest

Random Forest (RF) is a classifier that consists of multiple decision trees. A decision tree is made up of nodes and branches. At each node the dataset is split based on the value of some attribute that is selected so that the instances of different classes are predominantly moved to different branches. Classification starts at the root node and is performed by passing the instances along the tree to leaf nodes. To introduce diversity between the trees of a random forest, a subset of all attributes is randomly selected to take decisions at each node of each tree. The class probability of an instance is estimated considering results of all trees. We here developed RF models with 500 trees using the randomForest package of R. Thus, for a test set instance the class probability was one of 500 numerical values in the range from 0 to 1.

### Model evaluation

For every MoA/T, 25 RF models were created, assigning 80% of compounds to the training set and 20% of compounds to the test set. In each model, the seed for random number generator for training/test split corresponded to the model number to ensure reproducibility of modeling. The predictions from all models were aggregated to calculate Receiver Operating Characteristic (ROC) curve, which is plotted as the true positive rate versus the false positive rate at various discrimination threshold values. The area under the ROC curve (AUC) is a measure of the discriminatory power of a classifier, which is insensitive to class distributions and the costs of misclassifications; AUC = 1 indicates perfect classification, while AUC = 0.5 means that the classifier does not perform better than random guessing.

### Variable Importance in Random Forest models

In RF, each tree has an out-of-bag sample of data that is not used during construction of the tree. The prediction error on the out-of-bag portion of the data is calculated. Then, the values of the variables in the out-of-bag-sample are randomly shuffled, for one variable at a time, and predictions recalculated. The difference between the two values is used as a measure of importance of the variable in the given tree. Finally, the importance in the whole RF is calculated as the average over all trees.

## Results and Discussion

### Models for CMap datasets in three cell lines

In CMap dataset, the three cell lines with the highest number of annotated compounds were MCF7 (breast cancer, 2801 annotated compounds, PC3 (prostate cancer, 2775 annotated compounds), and A549 (lung cancer, 2319 annotated compounds). We developed Random Forest models for mechanisms of action and targets (MoA/Ts) shared by at least five compounds, which gave 444, 435, and 380 models for MCF7, PC3, and A549, respectively. In models for 17 MoA/Ts, the area under the ROC curve (AUC) exceeded 0.90, for 59 MoA/Ts the AUC exceeded 0.80, and for 121 MoA/Ts the AUC exceeded 0.70. The results for the best-predicted MoA/Ts are presented graphically in Figure 1 (for full results with number of active compounds in each model, AUC, and confidence intervals see Supplemental Table S1.)

**Figure 1.**
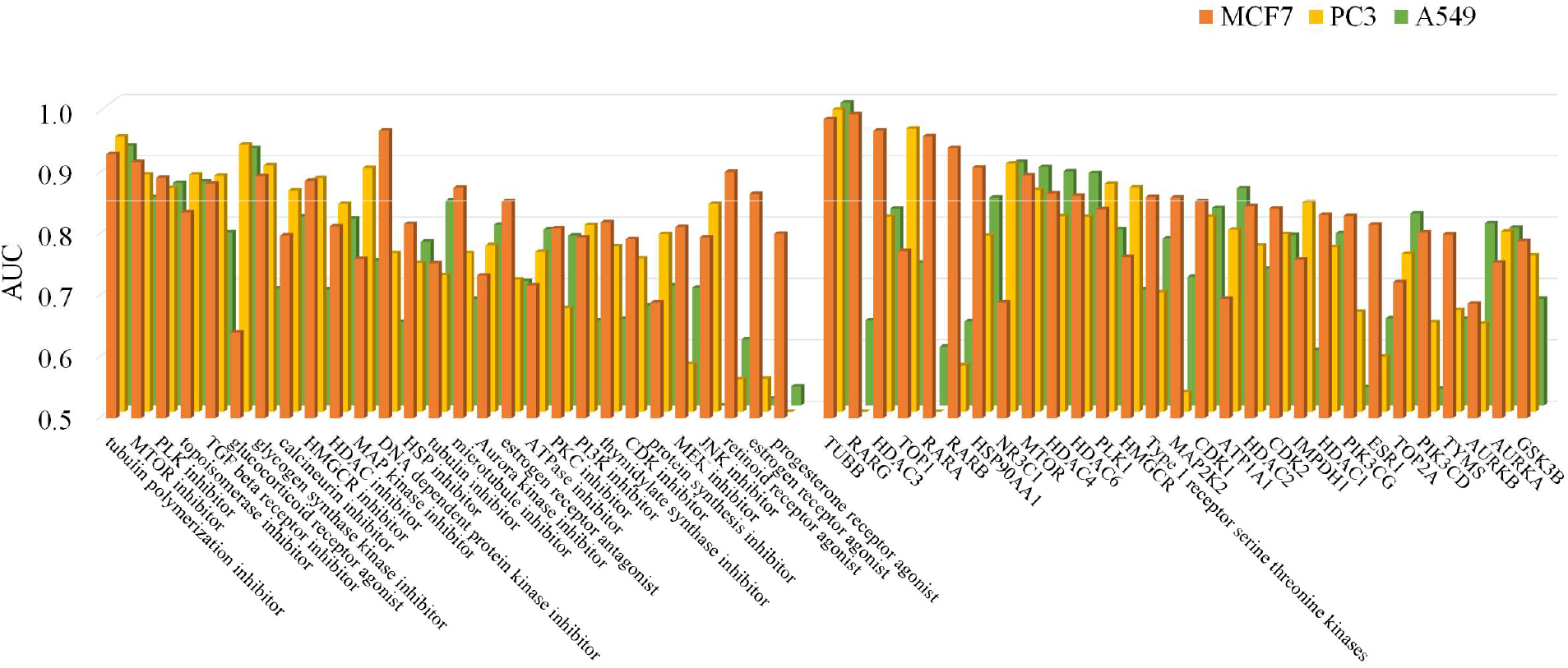
The areas under the ROC curve (AUC) for predicting MoA/Ts in cell lines MCF7, PC3, and A549. AUC is a measure of the discriminatory power of a model. AUC=1 indicates perfect predictions, i.e. complete separation of all class members from all non-members, whereas AUC=0.5 indicates predictions not better than random.

In the presentation of CMap dataset, the authors noted that only 15% of compounds produced highly similar transcriptional profiles across the entire panel of cell-lines suggesting that transcriptional response is cell dependent.^7^ For instance, it was found that glucocorticoid receptor antagonists shared similar profiles only in cell lines where the glucocorticoid receptor NR3C1 was highly expressed (i.e. A549, but not PC3 and MCF7). Our results confirm this finding for glucocorticoid receptor agonists, where the models for A549 and PC3 cell lines show much better predictive performance than the model for MCF7. Similarly, for glycogen synthase kinase inhibitors good models are obtained in MCF7 and PC3, but not in A549 cell line, but for estrogen receptor antagonists and agonists only in A549. For the three retinoid acid receptors RARA, RARB, and RARG, predictive models are obtained only in MCF7 cell line.

An overall comparison of the models also reveals some differences between the cell lines, the average AUC for the top-50 models in MCF7 cell line being 0.86, in PC3 cell line 0.83 and in A549 cell line 0.80. An overview of results for the broadest drug classes indicates that gene expression data is not suited for modeling of GPCR-targeted drugs (such as agonists and antagonists of dopamine, histamine, serotonin, and acetylcholine receptors). For these mechanisms of action, the models show AUC around 0.50, i.e., they do not perform better than random guesses. In contrast, an overall model for kinase inhibitors (that constitute about 10% of all dataset compounds) possesses predictive performance of AUC = 0.70 in MCF7 cell line and 0.71 in PC3.

### Models for CMap dataset in MCF7 cell line with gene expression profiles generated after different perturbation times

In the above presented models we used compound profiles generated after 24 hours of treatment. However, for MCF7 cell line, for 2787 of 2801 compounds profiles were available also for 6-hour treatment. In a preliminary analysis, we compared these two sets of profiles, and found that only for 56% of landmark genes the variance in expression levels was higher in 24-hour profiles compared to 6-hour profiles. This observation suggested that the effects of perturbations could not necessarily be manifested best in 24-hour profiles for all MoA/Ts.

Hence, to identify eventual time-dependencies, we created models also with 6-hour profiles and models where both profiles were concatenated, thus giving 2×978 features per compound. The results for 51 MoA/Ts where at least one of three models showed AUC > 0.8 are presented graphically in Figure 2. Notably, for some MoAs, such as tubulin polymerization inhibitor, microtubule inhibitor, and thymidylate synthase inhibitor, AUCs are low in models using 6-hour profiles. For most of MoA/Ts the model performances are, however, comparable, and in some cases combining of the two profiles has further improved the performance. Among the 51 MoAs/Ts, the highest AUC is achieved in 10 models built on 6-hour profiles, 26 models built on 24-hour profiles, and 15 models that include both profiles.

**Figure 2.**
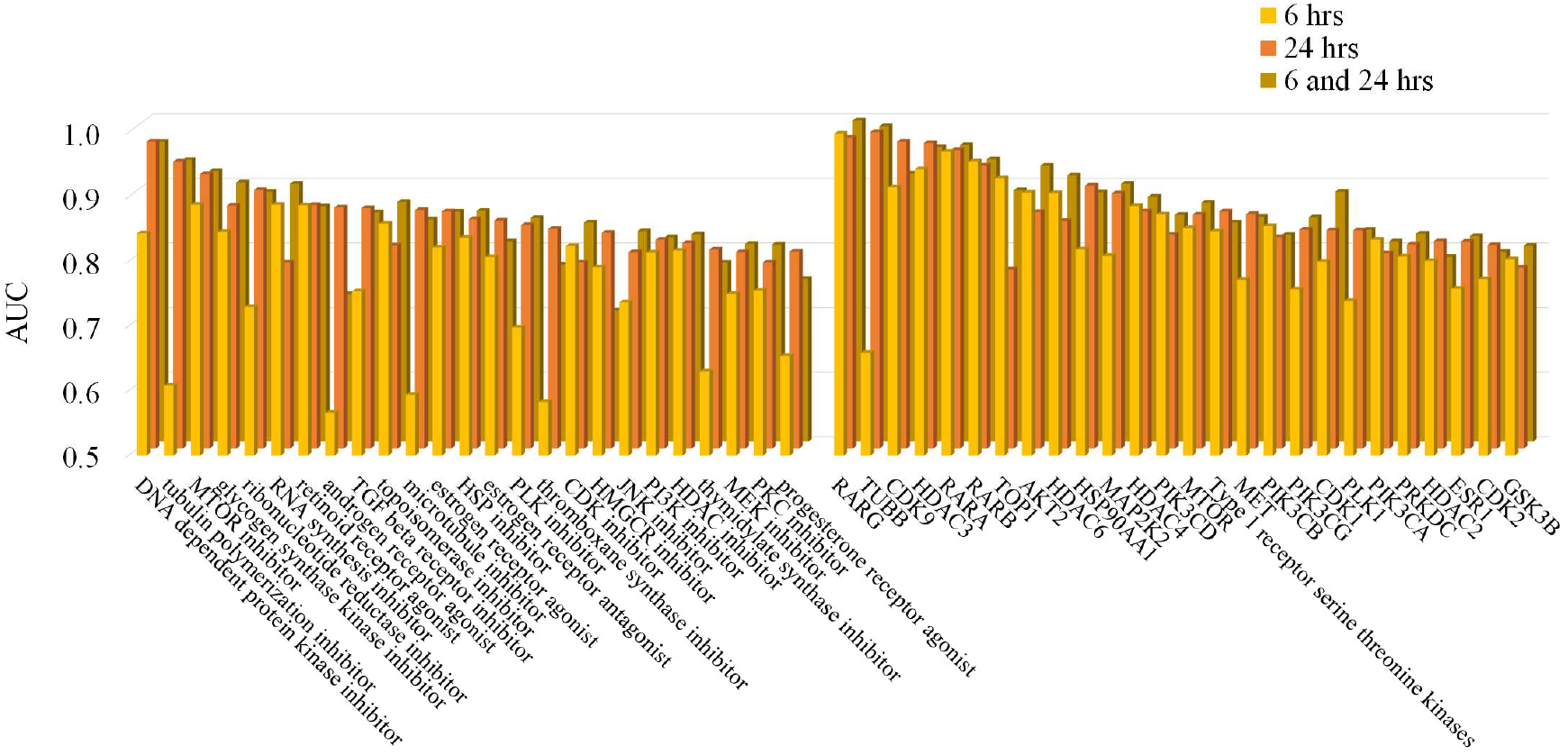
The areas under the ROC curve (AUC) in models for predicting MoA/Ts in MCF7 cell line using gene expression profiles generated after 6 hours and 24 hours of treatment, and a concatenation of the two profiles.

### Models for CMap/Cell Painting dataset

In the next step of the study we created models for a set of 1484 compounds that have been characterized in both gene expression and phenotypic profiling assays. For the sake of comparison, we also created models using structural descriptors of molecules, calculated by Chemistry Development Kit package of R (rcdk). These descriptors include a variety of topological, geometrical, charge based and constitutional descriptors.^18^

The results for MoA/Ts where AUC for either gene expression or phenotypic profiling based model exceeded 0.70 are presented in Figure 3 and Supplemental Table S2. In many cases, gene expression or phenotypic profiling models show comparable predictive performance. For some of the targets, however, only one of the two descriptions has produced a predictive model, suggesting that the two profiling approaches capture distinct aspects of cell response.

**Figure 3.**
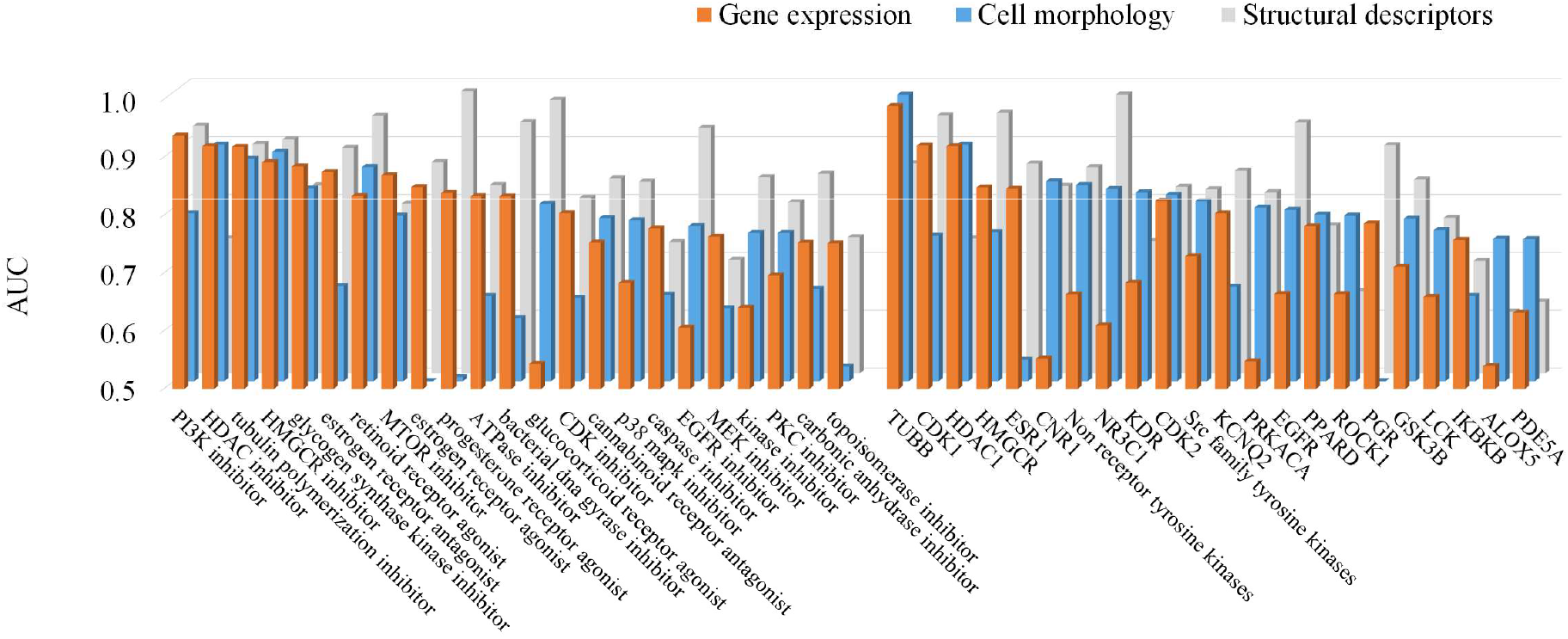
The areas under the ROC curve (AUC) for predicting MoA/Ts for models based on gene expression data, cell morphology data, and structural description of chemical compounds.

Comparisons of results for gene expression data based models in Figure 3 with the results of the corresponding models in Figures 1 and 2 reveal that for some MoA/Ts the modeling performance is inferior. This can be explained by a lower number of active compounds for these MoA/Ts in the intersection of CMap and Cell Painting datasets compared to CMap alone. For instance, the number of MTOR inhibitors has decreased from 18 to 5, resulting in decrease of AUC from 0.92 to 0.87. Similarly, the number of topoisomerase inhibitors has decreased from 21 to 7, resulting in change of AUC from 0.84 to 0.75.

Similarly as with gene expression data, phenotypic profiling data has not given any predictive models for agonists/antagonists of most GPCR classes. This is in contrast to models for several protein kinases and protein kinase groups (such as Non-receptor tyrosine kinases with AUC = 0.84 and Src family tyrosine kinases with AUC = 0.81) and nuclear receptors (e.g. glucocorticoid receptor NR3C1 with AUC = 0.83).

Our negative results for GPCRs are in agreement with findings of Rohban et. al.^19^ who estimated similarities of morphological profiles of pairs of compounds sharing the same MoA. For GPCR agonists/antagonists it was found that a very low fraction of the top most-similar profiles were profiles of compounds with the same MoA. Thus, for the four largest groups of compounds in the dataset, agonists and antagonists of dopamine and serotonin receptor, only 0 - 1% of top most-similar profiles belonged to another member of this group. (This can be compared to 2% for EGFR inhibitors, where we have obtained a predictive model with AUC = 0.77, 6% for PI3K inhibitors, where our model shows AUC = 0.79, and 18% for glucocorticoid receptor agonists, where our model shows AUC = 0.81).

In fact, for a multitude of MoA/Ts, the drug effect need not lead to profound morphological or transcriptional changes of cells. In profiling of 1600 known bioactive compounds by Cell Painting assay, Gustafsdottir et al. observed that only 13% of them could be deemed active, i.e. their profiles could be distinguished from the natural variation of profiles of untreated cells.^8^

### Interpretation of RF models

For RF models that use gene expression data, it is natural to ask if predictions are largely based on up/downregulation of a few key genes, or the whole panel of 978 landmark genes has been important for the modeling. Similarly, when models are built using cell morphology data, we can delve into which of the cell compartments, image channels, and Cell Painting feature groups have had the largest contributions.

Variable Importance (VI) for gene expression features in six of the models for drug targets are presented graphically in the left panel of Figure 4. In four of these models (TUBB, CDK1, HDAC1, and MAPK8) the five most important features accumulate 12% to 17% of the total sum of VI, whereas in two models (GSK3B and TOP2A) the fraction of VI accumulated by the five most important features is 10% and 6%, respectively. Notably, that for the former four models also the predictive performance is higher than for the latter two (cf. Supplemental Table S2). In all these models 50% of the sum of VI is accumulated by between 50 to 100 features. It can also be noted that many of the features have received VI values close to zero or negative values, which indicates that re-fitting of models after excluding these features would likely increase their predictive performances.

**Figure 4.**
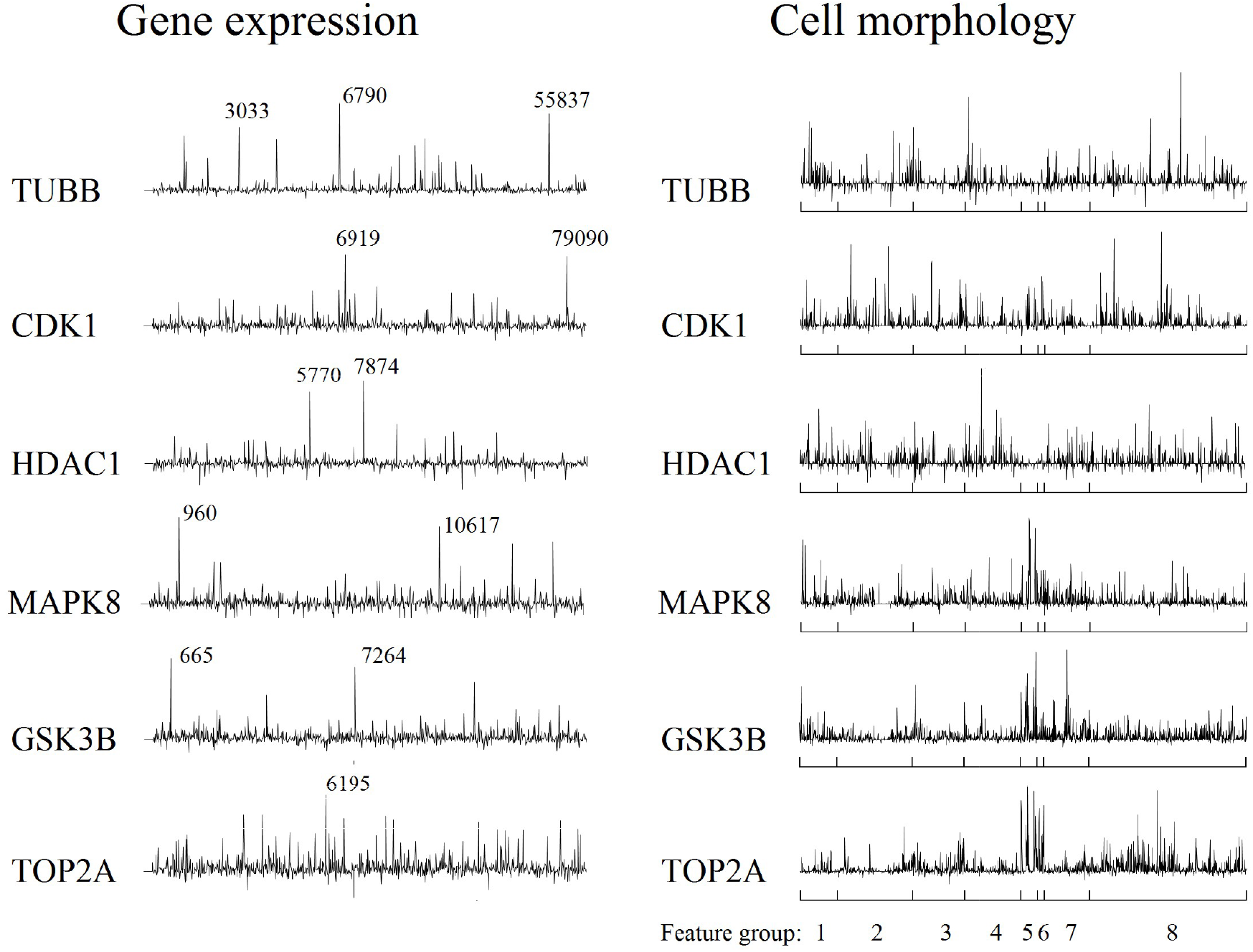
Variable Importances in models built using gene expression data (left panel) and cell morphology data (right panel). In the left panel, genes with highest VI are labeled by their NCBI Gene ID. In the right panel, indicated are eight CellProfiler feature groups: 1) AreaShape, 2) Correlation, 3) Granularity, 4) Intensity, 5) Location, 6) Neighbors, 7) RadialDistribution, and 8) Texture.

Some of the most important features in each model are labeled by their NCBI Gene ID. Inspection of data reveals that gene 6790 (AURKA), which appears to be among the most important in model for TUBB, is strongly upregulated in presence of all compounds that are associated with this target. This gene encodes for Aurora kinase A, which is involved in microtubule formation and stabilization. In contrast, gene 3033 (HADH) is strongly downregulated for all compounds and gene 55837 (EAPP) is downregulated for all compounds but one (mebendazole). Similar analysis can be performed for other models.

Variable Importances in corresponding models created with cell morphology features are presented in the right panel of Figure 4. The features are ordered according to Cell Painting feature groups (see Figure legend), and the pattern indicates that the most important features are distributed among all eight groups, although not evenly. Thus, models for TUBB and HDAC1 have not made much use of Location and Neighbors features, the VI values being close to 0. In contrast, these feature groups are important in models for MAPK8, GSK3B, and TOP2A, here less important being Correlation features. Interpreting the plot, one should take into account that the number of features in groups range from 21 (Neighbors) to 630 (Texture). Hence, even that the VI for individual Texture features in some of the models do not appear among the highest, the group as a whole accounts for 22% (for MAPK8) to 46% (for TUBB) of the sum of all VI.

### Models for Cell Painting dataset with different data pre-processing methods

In Cell Painting dataset, the majority of compounds are applied to cells eight times on different plates and locations on the plate and in this way eight morphological profiles per compound are obtained. The predominant concern is that phenotypic effects may be subtle and masked by systematic errors. Hence, a fraction of wells of each plate (typically, 1/6th) contains mock-treated (control) samples, which allows to reveal and compensate for the plate-to-plate variation and for the illumination heterogeneities at different locations on the plate.

In a preliminary study, we performed principal component analysis of morphological profiles of all control samples from 406 plates. The first principal component explained 25.6% of all variance and reflected primarily plate-wise deviations from the mean rather than intra-plate variation among the control samples (see Supplemental Figure S1).

To account for this, we attempted two pre-processing approaches: 1) centering of features on plate-to-plate basis by subtracting the mean value for the control samples on this plate and 2) centering and normalization by subtracting the mean and dividing by the standard deviation for the control samples. In the latter case, use of some features was problematic because the values for control samples were invariant or showed variance close to 0 for part of the plates.

Thereafter we described the compounds by either the mean values or the median values from the eight feature sets. In the latter case, the three “weakest” and the three “strongest” changes in cell morphology are not considered (note that some features became invariant when the median was selected instead of the mean).

Thus, four models could be built for each of the 262 MoA/Ts. Overall, the predictive performances of models were quite similar for most of MoA/Ts, the standard deviation calculated from the four AUC values being 0.033. However, larger deviations were observed for MoA/Ts with few active compounds. The dependency of modeling robustness on the number of active compounds is displayed graphically in Supplemental Figure S2.

The average AUC for top-50 best predicted MoA/Ts ranged from 0.748 (in models using mean values of centered data) to 0.760 (in models using mean values of centered and normalized data). From these consistent results we may conclude that modeling performance is not sensitive to the systematic errors that may occur during assaying large-scale datasets, provided that the compounds are applied to cells multiple times. Further studies are required to investigate if several Cell Painting datasets generated by different research groups and/or using several cell lines can be merged together to obtain a universal morphological description for any number of assayed compounds.

To complete the discussion, it should be noted that calculation of morphological features by CellProfiler is not mandatory for analysis of cell imaging data. Use of raw images as inputs to pre-trained convolutional neural networks has in fact shown to give better predictions in some studies.^13,20^

## Conclusion

Gene expression and phenotypic profiling of chemical compounds yield thousands of features that capture broad ranges of cellular responses to perturbation. These features can be considered as an alternative to structural descriptors of compounds in QSAR-like modeling. In this study we compared performances of gene expression profiles and phenotypic profiles in predicting compound mechanisms of action and targets (MoA/Ts). We identified MoA/Ts, for which both profiles gave predictive models, comparable to or better than structural descriptor based model. We also identified MoA/Ts, for which the performances of the two models were notably different, indicating that the two profiles have distinct information contents. This finding supports the idea of integrated approaches for predicting drug efficacy and safety in preclinical drug discovery.

For many MoA/Ts, the best model was obtained using structural descriptors. Interpreting this, one should take into account that any QSAR model has an applicability domain, limited by the structural diversity of the active compounds. For MoA/Ts with few active compounds, the model predictions cannot be extrapolated for the whole drug-like structure space. In contrast, the applicability domain of profiling-based description is limited by requirement for detectable cellular response but not by structural properties of the assayed compounds.

We also found that the prediction accuracy using gene expression features can be cell line specific, influenced by perturbation time, and proportional to the size of the dataset. Similar findings can be expected for phenotyping profiling data.

To our knowledge, our modeled gene expression and phenotypic profiling datasets from Broad Institute with 20K and 30K assayed compounds, respectively, are the largest publicly available at the present time. Further studies are warranted to confirm reproducibility of profiling methods and investigate if datasets generated by different research groups and/or using several cell lines can be merged together to obtain comparable quantitative descriptions for any number of assayed compounds.

