## Supplemental Material for "Evaluation of Gene Expression and Phenotypic Profiling Data as Quantitative Descriptors for Predicting Drug Targets and Mechanisms of Action"

**Table S1:** Results of modeling for CMap data set in cell lines MCF7, PC3, and A549, for mechanisms of action/targets (MoA/T), where in at least one of the three cell lines a predictive model was obtained with AUC > 0.7. Shown is the number of active compounds for the given MoA/T, AUC, and 95% confidence interval.

| **Cell line** | **MCF7** | | | **PC3** | | | **A549** | | |
|---|---|---|---|---|---|---|---|---|---|
| **MoA/T** | **No** | **AUC** | **95% CI** | **No** | **AUC** | **95% CI** | **No** | **AUC** | **95% CI** |
| Abl kinase inhibitor | 8 | 0.581 | 0.496 - 0.667 | 7 | 0.534 | 0.440 - 0.628 | 7 | 0.725 | 0.644 - 0.806 |
| adenosine receptor agonist | 12 | 0.410 | 0.347 - 0.472 | 12 | 0.704 | 0.633 - 0.776 | 11 | 0.631 | 0.570 - 0.691 |
| AKT inhibitor | 9 | 0.711 | 0.643 - 0.780 | 7 | 0.590 | 0.504 - 0.677 | 6 | 0.592 | 0.509 - 0.674 |
| AKT2 | 5 | 0.858 | 0.784 - 0.932 | 3 | | | 3 | | |
| androgen receptor agonist | 5 | 0.844 | 0.777 - 0.911 | 5 | 0.500 | 0.398 - 0.602 | 4 | | |
| antiarrhythmic | 5 | 0.719 | 0.606 - 0.831 | 5 | 0.688 | 0.579 - 0.798 | 5 | 0.476 | 0.381 - 0.570 |
| antioxidant | 11 | 0.460 | 0.402 - 0.519 | 11 | 0.582 | 0.509 - 0.656 | 6 | 0.730 | 0.657 - 0.802 |
| ATP1A1 | 15 | 0.695 | 0.621 - 0.770 | 17 | 0.797 | 0.738 - 0.856 | 13 | 0.854 | 0.793 - 0.916 |
| ATPase inhibitor | 24 | 0.717 | 0.665 - 0.769 | 26 | 0.761 | 0.711 - 0.810 | 24 | 0.787 | 0.732 - 0.841 |
| AURKA | 14 | 0.754 | 0.699 - 0.808 | 14 | 0.794 | 0.741 - 0.846 | 14 | 0.790 | 0.733 - 0.846 |
| AURKB | 12 | 0.687 | 0.623 - 0.752 | 13 | 0.644 | 0.563 - 0.726 | 12 | 0.797 | 0.746 - 0.847 |
| Aurora kinase inhibitor | 15 | 0.733 | 0.675 - 0.790 | 16 | 0.772 | 0.717 - 0.826 | 15 | 0.795 | 0.742 - 0.847 |
| Bcr-Abl kinase inhibitor | 7 | 0.629 | 0.541 - 0.717 | 6 | 0.576 | 0.474 - 0.679 | 6 | 0.770 | 0.685 - 0.855 |
| CA14 | 9 | 0.586 | 0.498 - 0.674 | 9 | 0.703 | 0.639 - 0.767 | 6 | 0.668 | 0.604 - 0.732 |
| CACNA1B | 7 | 0.515 | 0.427 - 0.602 | 7 | 0.571 | 0.486 - 0.656 | 6 | 0.712 | 0.659 - 0.764 |
| calcineurin inhibitor | 5 | 0.798 | 0.704 - 0.892 | 5 | 0.861 | 0.785 - 0.936 | 5 | 0.808 | 0.702 - 0.913 |
| CDK inhibitor | 24 | 0.792 | 0.741 - 0.844 | 22 | 0.750 | 0.690 - 0.811 | 21 | 0.663 | 0.598 - 0.728 |
| CDK1 | 16 | 0.854 | 0.803 - 0.905 | 15 | 0.818 | 0.762 - 0.875 | 14 | 0.822 | 0.764 - 0.880 |
| CDK2 | 18 | 0.842 | 0.793 - 0.891 | 18 | 0.790 | 0.723 - 0.858 | 18 | 0.778 | 0.723 - 0.833 |
| CDK9 | 5 | 0.951 | 0.906 - 0.997 | 4 | | | 4 | | |
| CFTR | 8 | 0.459 | 0.387 - 0.531 | 8 | 0.506 | 0.435 - 0.576 | 7 | 0.732 | 0.661 - 0.804 |
| DHFR | 6 | 0.636 | 0.519 - 0.753 | 6 | 0.781 | 0.695 - 0.866 | 3 | | |
| dihydrofolate reductase inhibitor | 5 | 0.713 | 0.595 - 0.831 | 5 | 0.792 | 0.691 - 0.893 | 3 | | |
| DNA dependent protein kinase inhibitor | 5 | 0.969 | 0.953 - 0.986 | 5 | 0.759 | 0.645 - 0.874 | 5 | 0.636 | 0.509 - 0.763 |
| DNMT1 | 7 | 0.565 | 0.490 - 0.639 | 6 | 0.499 | 0.418 - 0.579 | 6 | 0.713 | 0.650 - 0.775 |
| Erb-b2 receptor tyrosine kinases | 45 | 0.646 | 0.614 - 0.678 | 45 | 0.703 | 0.666 - 0.740 | 42 | 0.659 | 0.625 - 0.693 |
| ESR1 | 39 | 0.816 | 0.783 - 0.848 | 39 | 0.590 | 0.549 - 0.631 | 34 | 0.642 | 0.597 - 0.687 |
| ESR2 | 23 | 0.736 | 0.684 - 0.788 | 23 | 0.505 | 0.453 - 0.556 | 21 | 0.543 | 0.480 - 0.605 |

| | | | | | | | | | |
|---|---|---|---|---|---|---|---|---|---|
| estrogen receptor agonist | 29 | 0.866 | 0.834 - 0.898 | 29 | 0.554 | 0.506 - 0.601 | 26 | 0.511 | 0.466 - 0.556 |
| estrogen receptor antagonist | 11 | 0.854 | 0.778 - 0.931 | 11 | 0.716 | 0.66 - 0.773 | 11 | 0.703 | 0.622 - 0.785 |
| FLT4 | 15 | 0.717 | 0.655 - 0.778 | 13 | 0.627 | 0.564 - 0.69 | 12 | 0.679 | 0.610 - 0.748 |
| glucocorticoid receptor agonist | 49 | 0.640 | 0.609 - 0.672 | 49 | 0.936 | 0.916 - 0.956 | 40 | 0.920 | 0.889 - 0.950 |
| glutamate receptor modulator | 9 | 0.578 | 0.496 - 0.660 | 9 | 0.370 | 0.312 - 0.428 | 8 | 0.810 | 0.746 - 0.875 |
| glycogen synthase kinase inhibitor | 12 | 0.895 | 0.851 - 0.940 | 11 | 0.902 | 0.869 - 0.936 | 11 | 0.691 | 0.609 - 0.773 |
| growth factor receptor inhibitor | 8 | 0.724 | 0.647 - 0.800 | 5 | 0.656 | 0.543 - 0.769 | 5 | 0.572 | 0.479 - 0.664 |
| GSK3B | 22 | 0.789 | 0.745 - 0.833 | 20 | 0.755 | 0.700 - 0.810 | 20 | 0.674 | 0.617 - 0.732 |
| HDAC inhibitor | 32 | 0.813 | 0.762 - 0.864 | 32 | 0.839 | 0.798 - 0.879 | 29 | 0.805 | 0.754 - 0.855 |
| HDAC1 | 23 | 0.832 | 0.782 - 0.882 | 23 | 0.769 | 0.711 - 0.826 | 20 | 0.781 | 0.718 - 0.844 |
| HDAC2 | 17 | 0.846 | 0.790 - 0.902 | 17 | 0.771 | 0.701 - 0.841 | 16 | 0.723 | 0.648 - 0.799 |
| HDAC3 | 15 | 0.969 | 0.949 - 0.988 | 15 | 0.818 | 0.749 - 0.887 | 14 | 0.821 | 0.753 - 0.890 |
| HDAC4 | 9 | 0.867 | 0.802 - 0.933 | 9 | 0.819 | 0.734 - 0.903 | 8 | 0.882 | 0.816 - 0.948 |
| HDAC6 | 18 | 0.863 | 0.814 - 0.913 | 18 | 0.818 | 0.759 - 0.877 | 15 | 0.879 | 0.827 - 0.931 |
| histone lysine methyltransferase inhibitor | 4 | | | 5 | 0.675 | 0.565 - 0.785 | 6 | 0.731 | 0.631 - 0.832 |
| HMGCR | 12 | 0.763 | 0.685 - 0.842 | 12 | 0.866 | 0.806 - 0.926 | 9 | 0.689 | 0.605 - 0.773 |
| HMGCR inhibitor | 11 | 0.887 | 0.838 - 0.936 | 11 | 0.881 | 0.824 - 0.938 | 9 | 0.689 | 0.605 - 0.773 |
| HSD11B1 | 5 | 0.356 | 0.287 - 0.424 | 5 | 0.608 | 0.515 - 0.700 | 5 | 0.700 | 0.645 - 0.754 |
| HSP inhibitor | 10 | 0.817 | 0.752 - 0.881 | 12 | 0.743 | 0.661 - 0.826 | 10 | 0.767 | 0.693 - 0.841 |
| HSP90AA1 | 7 | 0.909 | 0.858 - 0.960 | 8 | 0.787 | 0.701 - 0.873 | 7 | 0.839 | 0.761 - 0.918 |
| IKK inhibitor | 8 | 0.625 | 0.542 - 0.709 | 8 | 0.610 | 0.526 - 0.694 | 8 | 0.716 | 0.641 - 0.791 |
| IMPDH1 | 6 | 0.759 | 0.676 - 0.842 | 6 | 0.842 | 0.765 - 0.919 | 5 | 0.590 | 0.469 - 0.712 |
| inositol monophosphatase inhibitor | 5 | 0.715 | 0.593 - 0.838 | 5 | 0.619 | 0.496 - 0.742 | 5 | 0.727 | 0.628 - 0.827 |
| insulin sensitizer | 5 | 0.665 | 0.601 - 0.729 | 5 | 0.893 | 0.796 - 0.990 | 4 | | |
| JNK inhibitor | 5 | 0.795 | 0.694 - 0.896 | 5 | 0.839 | 0.749 - 0.929 | 5 | 0.430 | 0.329 - 0.531 |
| KCNQ2 | 6 | 0.720 | 0.665 - 0.775 | 6 | 0.603 | 0.500 - 0.705 | 6 | 0.602 | 0.534 - 0.670 |
| KDR | 32 | 0.734 | 0.695 - 0.774 | 28 | 0.608 | 0.558 - 0.659 | 26 | 0.645 | 0.599 - 0.691 |
| kinase inhibitor | 248 | 0.703 | 0.678 - 0.728 | 230 | 0.708 | 0.688 - 0.738 | 225 | 0.669 | 0.648 - 0.700 |
| KIT inhibitor | 14 | 0.677 | 0.607 - 0.747 | 12 | 0.696 | 0.625 - 0.766 | 11 | 0.731 | 0.663 - 0.800 |
| LCK | 18 | 0.659 | 0.601 - 0.718 | 16 | 0.650 | 0.586 - 0.713 | 16 | 0.716 | 0.661 - 0.771 |
| MAP kinase inhibitor | 6 | 0.760 | 0.672 - 0.848 | 6 | 0.898 | 0.825 - 0.971 | 6 | 0.736 | 0.628 - 0.844 |
| MAP2K1 | 14 | 0.748 | 0.682 - 0.813 | 13 | 0.432 | 0.368 - 0.496 | 12 | 0.577 | 0.496 - 0.657 |
| MAP2K2 | 7 | 0.860 | 0.798 - 0.922 | 6 | 0.532 | 0.424 - 0.639 | 6 | 0.710 | 0.621 - 0.800 |
| MDM2 | 5 | 0.700 | 0.581 - 0.819 | 5 | 0.584 | 0.503 - 0.666 | 5 | 0.543 | 0.441 - 0.645 |
| MEK inhibitor | 16 | 0.812 | 0.746 - 0.879 | 13 | 0.578 | 0.513 - 0.642 | 12 | 0.692 | 0.624 - 0.760 |
| MET | 12 | 0.902 | 0.863 - 0.941 | 4 | | | 4 | | |
| microtubule inhibitor | 8 | 0.876 | 0.825 - 0.927 | 8 | 0.759 | 0.667 - 0.852 | 7 | 0.674 | 0.588 - 0.761 |
| MMP13 | 6 | 0.741 | 0.652 - 0.830 | 6 | 0.563 | 0.465 - 0.662 | 6 | 0.618 | 0.552 - 0.684 |
| MTOR | 18 | 0.896 | 0.860 - 0.931 | 15 | 0.862 | 0.808 - 0.916 | 16 | 0.889 | 0.848 - 0.929 |
| MTOR inhibitor | 18 | 0.918 | 0.893 - 0.943 | 16 | 0.887 | 0.837 - 0.937 | 17 | 0.840 | 0.788 - 0.892 |
| neuropeptide receptor antagonist | 5 | 0.436 | 0.343 - 0.528 | 5 | 0.751 | 0.637 - 0.866 | 4 | | |
| nitric oxide production inhibitor | 6 | 0.574 | 0.474 - 0.674 | 7 | 0.424 | 0.349 - 0.499 | 5 | 0.718 | 0.615 - 0.821 |
| NR1I3 | 5 | 0.525 | 0.429 - 0.620 | 5 | 0.378 | 0.309 - 0.447 | 5 | 0.704 | 0.639 - 0.770 |
| NR3C1 | 52 | 0.689 | 0.660 - 0.718 | 52 | 0.905 | 0.881 - 0.929 | 46 | 0.897 | 0.864 - 0.929 |
| other antibiotic | 6 | 0.702 | 0.608 - 0.795 | 6 | 0.753 | 0.660 - 0.846 | 2 | | |
| p38 MAPK inhibitor | 16 | 0.646 | 0.586 - 0.706 | 16 | 0.719 | 0.662 - 0.777 | 15 | 0.558 | 0.484 - 0.632 |
| PDE2A | 5 | 0.357 | 0.289 - 0.426 | 5 | 0.745 | 0.639 - 0.851 | 5 | 0.717 | 0.671 - 0.762 |
| PDPK1 | 5 | 0.730 | 0.649 - 0.811 | 5 | 0.489 | 0.374 - 0.605 | 5 | 0.708 | 0.605 - 0.810 |
| PI3K inhibitor | 22 | 0.795 | 0.741 - 0.850 | 18 | 0.805 | 0.746 - 0.864 | 17 | 0.639 | 0.580 - 0.698 |
| PIK3CA | 12 | 0.787 | 0.711 - 0.863 | 10 | 0.652 | 0.554 - 0.749 | 10 | 0.637 | 0.570 - 0.704 |
| PIK3CB | 9 | 0.801 | 0.721 - 0.882 | 8 | 0.690 | 0.596 - 0.785 | 8 | 0.510 | 0.431 - 0.588 |
| PIK3CD | 11 | 0.803 | 0.730 - 0.877 | 10 | 0.646 | 0.551 - 0.742 | 10 | 0.527 | 0.452 - 0.602 |
| PIK3CG | 16 | 0.830 | 0.779 - 0.882 | 13 | 0.663 | 0.576 - 0.750 | 12 | 0.530 | 0.474 - 0.587 |
| PKC inhibitor | 13 | 0.810 | 0.689 - 0.815 | 13 | 0.669 | 0.591 - 0.747 | 12 | 0.777 | 0.716 - 0.894 |
| PLK inhibitor | 7 | 0.892 | 0.842 - 0.941 | 6 | 0.865 | 0.793 - 0.937 | 5 | 0.863 | 0.784 - 0.941 |
| PLK1 | 8 | 0.841 | 0.783 - 0.900 | 8 | 0.872 | 0.808 - 0.935 | 6 | 0.787 | 0.709 - 0.865 |
| PRKCB | 5 | 0.504 | 0.399 - 0.609 | 5 | 0.491 | 0.383 - 0.598 | 5 | 0.706 | 0.599 - 0.813 |
| PRKDC | 8 | 0.749 | 0.661 - 0.837 | 8 | 0.749 | 0.665 - 0.833 | 7 | 0.558 | 0.460 - 0.657 |
| progesterone receptor agonist | 20 | 0.801 | 0.755 - 0.846 | 21 | 0.480 | 0.433 - 0.528 | 16 | 0.531 | 0.466 - 0.597 |
| progesterone receptor antagonist | 5 | 0.550 | 0.448 - 0.652 | 5 | 0.794 | 0.696 - 0.892 | 3 | | |
| progestogen hormone | 5 | 0.749 | 0.648 - 0.850 | 5 | 0.609 | 0.518 - 0.700 | 4 | | |
| protein synthesis inhibitor | 17 | 0.689 | 0.619 - 0.760 | 19 | 0.790 | 0.736 - 0.844 | 16 | 0.696 | 0.614 - 0.778 |
| PTPN1 | 5 | 0.353 | 0.287 - 0.420 | 5 | 0.776 | 0.690 - 0.861 | 5 | 0.553 | 0.471 - 0.636 |
| RARA | 12 | 0.960 | 0.940 - 0.980 | 12 | 0.430 | 0.373 - 0.487 | 12 | 0.596 | 0.528 - 0.664 |

| | | | | | | | | | |
|---|---|---|---|---|---|---|---|---|---|
| RARB | 11 | 0.941 | 0.900 - 0.982 | 11 | 0.576 | 0.515 - 0.637 | 11 | 0.637 | 0.558 - 0.716 |
| RARG | 6 | 0.996 | 0.992 - 0.999 | 6 | 0.436 | 0.361 - 0.510 | 6 | 0.639 | 0.550 - 0.727 |
| Receptor tyrosine kinases other | 78 | 0.707 | 0.678 - 0.735 | 67 | 0.638 | 0.607 - 0.668 | 65 | 0.691 | 0.664 - 0.719 |
| RET | 11 | 0.751 | 0.690 - 0.813 | 10 | 0.686 | 0.606 - 0.765 | 10 | 0.665 | 0.593 - 0.737 |
| retinoid receptor agonist | 15 | 0.902 | 0.858 - 0.946 | 15 | 0.553 | 0.502 - 0.605 | 15 | 0.608 | 0.541 - 0.674 |
| ribonucleotide reductase inhibitor | 5 | 0.747 | 0.623 - 0.870 | 5 | 0.683 | 0.562 - 0.804 | 2 | | |
| RNA synthesis inhibitor | 5 | 0.762 | 0.666 - 0.858 | 5 | 0.762 | 0.664 - 0.861 | 3 | | |
| RPL3 | 4 | | | 5 | 0.998 | 0.997 - 0.999 | 5 | 0.995 | 0.993 - 0.997 |
| RRM1 | 5 | 0.688 | 0.564 - 0.811 | 5 | 0.791 | 0.693 - 0.890 | 2 | | |
| sigma receptor antagonist | 5 | 0.626 | 0.510 - 0.742 | 5 | 0.736 | 0.617 - 0.855 | 5 | 0.726 | 0.683 - 0.769 |
| SRC | 12 | 0.724 | 0.657 - 0.792 | 10 | 0.737 | 0.655 - 0.819 | 10 | 0.764 | 0.697 - 0.832 |
| SRC family tyrosine kinases | 25 | 0.666 | 0.613 - 0.720 | 24 | 0.662 | 0.607 - 0.716 | 23 | 0.753 | 0.704 - 0.801 |
| TGF beta receptor inhibitor | 5 | 0.883 | 0.780 - 0.987 | 5 | 0.885 | 0.804 - 0.966 | 5 | 0.782 | 0.664 - 0.900 |
| TH | 5 | 0.470 | 0.378 - 0.563 | 5 | 0.708 | 0.605 - 0.811 | 2 | | |
| thromboxane synthase inhibitor | 5 | 0.807 | 0.725 - 0.889 | 5 | 0.629 | 0.526 - 0.732 | 4 | | |
| thymidylate synthase inhibitor | 7 | 0.820 | 0.749 - 0.890 | 7 | 0.770 | 0.682 - 0.858 | 5 | 0.641 | 0.537 - 0.744 |
| TK2 | 6 | 0.667 | 0.565 - 0.769 | 5 | 0.726 | 0.617 - 0.834 | 5 | 0.684 | 0.568 - 0.800 |
| TOP1 | 7 | 0.773 | 0.688 - 0.859 | 7 | 0.962 | 0.93 - 0.993 | 6 | 0.733 | 0.621 - 0.844 |
| TOP2A | 23 | 0.722 | 0.664 - 0.780 | 24 | 0.758 | 0.709 - 0.806 | 17 | 0.813 | 0.754 - 0.872 |
| topoisomerase inhibitor | 21 | 0.836 | 0.789 - 0.884 | 22 | 0.887 | 0.842 - 0.933 | 18 | 0.865 | 0.808 - 0.922 |
| TP53 | 5 | 0.702 | 0.593 - 0.811 | 5 | 0.491 | 0.384 - 0.599 | 4 | | |
| TRPM3 | 5 | 0.563 | 0.449 - 0.677 | 5 | 0.786 | 0.662 - 0.910 | 5 | 0.772 | 0.677 - 0.866 |
| TUBB | 16 | 0.988 | 0.984 - 0.992 | 16 | 0.993 | 0.990 - 0.996 | 14 | 0.994 | 0.991 - 0.996 |
| tubulin inhibitor | 6 | 0.753 | 0.654 - 0.852 | 6 | 0.723 | 0.627 - 0.82 | 5 | 0.834 | 0.742 - 0.925 |
| tubulin polymerization inhibitor | 13 | 0.931 | 0.893 - 0.969 | 13 | 0.949 | 0.921 - 0.977 | 12 | 0.924 | 0.875 - 0.973 |
| TYMS | 9 | 0.800 | 0.736 - 0.863 | 9 | 0.666 | 0.578 - 0.754 | 5 | 0.641 | 0.537 - 0.744 |
| Type 1 receptor serine threonine kinases | 6 | 0.861 | 0.782 - 0.940 | 6 | 0.695 | 0.593 - 0.796 | 6 | 0.772 | 0.708 - 0.836 |
| VDR | 8 | 0.798 | 0.722 - 0.873 | 8 | 0.680 | 0.586 - 0.775 | 6 | 0.495 | 0.404 - 0.586 |
| vitamin D receptor agonist | 8 | 0.798 | 0.722 - 0.873 | 8 | 0.680 | 0.586 - 0.775 | 6 | 0.495 | 0.404 - 0.586 |

**Table S2:** Results of modeling for CMap/Cell Painting data set for mechanisms of action/targets (MoA/T), where a predictive model with AUC > 0.7 was obtained using gene expression data and/or cell morphology data. Shown is the number of active compounds for the given MoA/T, AUC, and 95% confidence interval.

| **MoA/T** | | **Gene expression** | | **Cell morphology** | | **Structural descriptors** | |
|---|---|---|---|---|---|---|---|
| | **No** | **AUC** | **95% CI** | **AUC** | **95% CI** | **AUC** | **95% CI** |
| ALOX5 | 9 | 0.539 | 0.467 - 0.612 | 0.746 | 0.672 - 0.820 | 0.606 | 0.516 - 0.697 |
| angiotensin converting enzyme inhibitor | 6 | 0.635 | 0.529 - 0.740 | 0.735 | 0.652 - 0.818 | 0.837 | 0.754 - 0.919 |
| antiarrhythmic | 5 | 0.704 | 0.573 - 0.835 | 0.624 | 0.505 - 0.744 | 0.798 | 0.683 - 0.912 |
| ATPase inhibitor | 7 | 0.833 | 0.752 - 0.914 | 0.647 | 0.552 - 0.741 | 0.826 | 0.740 - 0.912 |
| bacterial DNA gyrase inhibitor | 10 | 0.833 | 0.767 - 0.898 | 0.609 | 0.525 - 0.694 | 0.934 | 0.882 - 0.986 |
| BCHE | 5 | 0.663 | 0.588 - 0.738 | 0.739 | 0.656 - 0.823 | 0.849 | 0.768 - 0.929 |
| CA1 | 25 | 0.567 | 0.505 - 0.629 | 0.715 | 0.665 - 0.764 | 0.860 | 0.821 - 0.900 |
| cannabinoid receptor antagonist | 6 | 0.753 | 0.675 - 0.830 | 0.782 | 0.715 - 0.849 | 0.837 | 0.761 - 0.913 |
| carbonic anhydrase inhibitor | 9 | 0.752 | 0.688 - 0.817 | 0.659 | 0.586 - 0.732 | 0.845 | 0.773 - 0.918 |
| caspase inhibitor | 5 | 0.777 | 0.684 - 0.871 | 0.649 | 0.519 - 0.780 | 0.727 | 0.616 - 0.838 |
| CDK inhibitor | 12 | 0.804 | 0.733 - 0.874 | 0.644 | 0.548 - 0.740 | 0.804 | 0.726 - 0.881 |
| CDK1 | 5 | 0.920 | 0.868 - 0.973 | 0.751 | 0.631 - 0.871 | 0.946 | 0.899 - 0.993 |
| CDK2 | 7 | 0.825 | 0.758 - 0.892 | 0.822 | 0.747 - 0.897 | 0.823 | 0.733 - 0.913 |
| CHEK1 | 7 | 0.625 | 0.541 - 0.709 | 0.708 | 0.615 - 0.802 | 0.806 | 0.742 - 0.870 |
| chloride channel blocker | 6 | 0.412 | 0.333 - 0.491 | 0.732 | 0.670 - 0.794 | 0.818 | 0.727 - 0.909 |
| CNR1 | 13 | 0.553 | 0.477 - 0.628 | 0.846 | 0.785 - 0.906 | 0.824 | 0.765 - 0.883 |
| CNR2 | 13 | 0.726 | 0.657 - 0.796 | 0.535 | 0.469 - 0.601 | 0.810 | 0.755 - 0.865 |
| EGFR | 20 | 0.664 | 0.620 - 0.708 | 0.796 | 0.752 - 0.839 | 0.934 | 0.906 - 0.961 |
| EGFR inhibitor | 23 | 0.606 | 0.560 - 0.652 | 0.768 | 0.722 - 0.814 | 0.925 | 0.908 - 0.941 |

| | | | | | | | |
|---|---|---|---|---|---|---|---|
| ERBB2 | 5 | 0.681 | 0.644 - 0.717 | 0.744 | 0.698 - 0.790 | 0.942 | 0.916 - 0.968 |
| ESR1 | 17 | 0.846 | 0.813 - 0.878 | 0.537 | 0.474 - 0.600 | 0.863 | 0.815 - 0.910 |
| ESR2 | 13 | 0.703 | 0.637 - 0.769 | 0.539 | 0.476 - 0.603 | 0.832 | 0.777 - 0.887 |
| estrogen receptor agonist | 11 | 0.849 | 0.793 - 0.904 | 0.492 | 0.412 - 0.571 | 0.865 | 0.813 - 0.917 |
| estrogen receptor antagonist | 6 | 0.874 | 0.809 - 0.940 | 0.664 | 0.572 - 0.756 | 0.890 | 0.833 - 0.947 |
| glucocorticoid receptor agonist | 14 | 0.543 | 0.476 - 0.611 | 0.806 | 0.746 - 0.867 | 0.973 | 0.946 - 0.999 |
| glycogen synthase kinase inhibitor | 6 | 0.884 | 0.836 - 0.933 | 0.833 | 0.771 - 0.896 | 0.825 | 0.743 - 0.908 |
| GSK3B | 11 | 0.711 | 0.639 - 0.783 | 0.781 | 0.713 - 0.848 | 0.835 | 0.789 - 0.881 |
| HDAC inhibitor | 5 | 0.919 | 0.837 - 1.000 | 0.909 | 0.826 - 0.992 | 0.734 | 0.635 - 0.832 |
| HDAC1 | 5 | 0.919 | 0.837 - 1.000 | 0.909 | 0.826 - 0.992 | 0.734 | 0.635 - 0.832 |
| HMGCR | 5 | 0.848 | 0.753 - 0.944 | 0.757 | 0.634 - 0.879 | 0.951 | 0.883 - 1.000 |
| HMGCR inhibitor | 5 | 0.892 | 0.837 - 0.946 | 0.896 | 0.849 - 0.943 | 0.904 | 0.817 - 0.992 |
| IKBKB | 6 | 0.757 | 0.648 - 0.867 | 0.647 | 0.554 - 0.74 | 0.695 | 0.611 - 0.779 |
| KCNQ2 | 5 | 0.803 | 0.769 - 0.838 | 0.663 | 0.554 - 0.771 | 0.851 | 0.747 - 0.954 |
| KDR | 7 | 0.683 | 0.584 - 0.782 | 0.826 | 0.755 - 0.897 | 0.729 | 0.639 - 0.819 |
| kinase inhibitor | 101 | 0.640 | 0.614 - 0.667 | 0.747 | 0.721 - 0.772 | 0.839 | 0.818 - 0.860 |
| LCK | 12 | 0.659 | 0.585 - 0.732 | 0.761 | 0.692 - 0.829 | 0.769 | 0.714 - 0.824 |
| lipoxygenase inhibitor | 6 | 0.713 | 0.636 - 0.790 | 0.709 | 0.658 - 0.760 | 0.719 | 0.630 - 0.809 |
| MAPK1 | 19 | 0.703 | 0.642 - 0.763 | 0.671 | 0.600 - 0.741 | 0.775 | 0.722 - 0.829 |
| MAPK8 | 8 | 0.713 | 0.626 - 0.800 | 0.694 | 0.619 - 0.769 | 0.751 | 0.681 - 0.822 |
| MEK inhibitor | 6 | 0.763 | 0.659 - 0.867 | 0.626 | 0.510 - 0.741 | 0.696 | 0.589 - 0.804 |
| MTOR inhibitor | 5 | 0.869 | 0.796 - 0.941 | 0.786 | 0.684 - 0.888 | 0.794 | 0.686 - 0.901 |
| Non-receptor tyrosine kinases | 24 | 0.663 | 0.606 - 0.721 | 0.839 | 0.805 - 0.873 | 0.857 | 0.824 - 0.889 |
| norepinephrine reuptake inhibitor | 5 | 0.535 | 0.424 - 0.646 | 0.706 | 0.641 - 0.772 | 0.716 | 0.609 - 0.822 |
| NR1I2 | 5 | 0.710 | 0.622 - 0.799 | 0.514 | 0.421 - 0.606 | 0.569 | 0.463 - 0.675 |
| NR3C1 | 18 | 0.610 | 0.554 - 0.665 | 0.832 | 0.788 - 0.877 | 0.982 | 0.967 - 0.997 |
| p38 MAPK inhibitor | 10 | 0.683 | 0.605 - 0.761 | 0.778 | 0.706 - 0.850 | 0.832 | 0.762 - 0.901 |
| PDE5A | 12 | 0.631 | 0.576 - 0.687 | 0.745 | 0.685 - 0.805 | 0.625 | 0.541 - 0.708 |
| PGR | 11 | 0.786 | 0.729 - 0.843 | 0.486 | 0.435 - 0.536 | 0.895 | 0.846 - 0.943 |
| PI3K inhibitor | 5 | 0.938 | 0.910 - 0.965 | 0.790 | 0.682 - 0.898 | 0.928 | 0.857 - 0.998 |
| PKC inhibitor | 7 | 0.696 | 0.602 - 0.790 | 0.756 | 0.682 - 0.830 | 0.796 | 0.723 - 0.869 |
| PPARD | 5 | 0.781 | 0.711 - 0.851 | 0.788 | 0.749 - 0.828 | 0.756 | 0.671 - 0.841 |
| PRKACA | 6 | 0.548 | 0.472 - 0.624 | 0.800 | 0.690 - 0.911 | 0.813 | 0.714 - 0.912 |
| progesterone receptor agonist | 6 | 0.839 | 0.776 - 0.901 | 0.507 | 0.424 - 0.589 | 0.988 | 0.980 - 0.995 |
| Receptor tyrosine kinases | 43 | 0.583 | 0.543 - 0.622 | 0.700 | 0.661 - 0.740 | 0.854 | 0.823 - 0.885 |
| retinoid receptor agonist | 7 | 0.833 | 0.745 - 0.922 | 0.870 | 0.787 - 0.954 | 0.945 | 0.910 - 0.980 |
| ROCK1 | 10 | 0.664 | 0.590 - 0.738 | 0.786 | 0.725 - 0.847 | 0.643 | 0.574 - 0.711 |
| Src family tyrosine kinases | 15 | 0.729 | 0.662 - 0.795 | 0.811 | 0.755 - 0.867 | 0.818 | 0.772 - 0.864 |
| SRC inhibitor | 7 | 0.729 | 0.660 - 0.797 | 0.661 | 0.574 - 0.747 | 0.722 | 0.646 - 0.798 |
| TNF | 8 | 0.712 | 0.642 - 0.782 | 0.493 | 0.407 - 0.580 | 0.523 | 0.448 - 0.598 |
| TOP2A | 12 | 0.669 | 0.604 - 0.735 | 0.705 | 0.629 - 0.781 | 0.899 | 0.847 - 0.952 |
| topoisomerase inhibitor | 7 | 0.752 | 0.687 - 0.816 | 0.525 | 0.432 - 0.618 | 0.735 | 0.648 - 0.823 |
| transferase inhibitor | 6 | 0.733 | 0.639 - 0.827 | 0.489 | 0.392 - 0.586 | 0.787 | 0.693 - 0.880 |
| TUBB | 7 | 0.989 | 0.984 - 0.993 | 0.995 | 0.993 - 0.997 | 0.863 | 0.797 - 0.929 |
| tubulin polymerization inhibitor | 7 | 0.918 | 0.867 - 0.969 | 0.884 | 0.810 - 0.958 | 0.897 | 0.822 - 0.971 |
| tyrosine kinase inhibitor | 69 | 0.619 | 0.587 - 0.651 | 0.747 | 0.716 - 0.777 | 0.872 | 0.849 - 0.895 |
| VEGFR inhibitor | 7 | 0.592 | 0.501 - 0.682 | 0.740 | 0.639 - 0.842 | 0.807 | 0.741 - 0.872 |

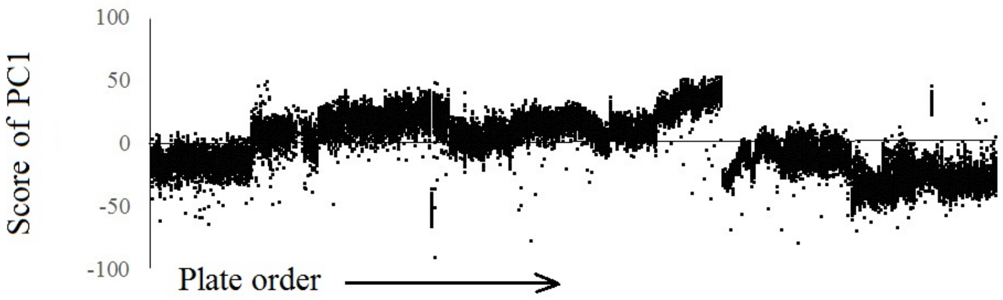


**Figure S1.** The first component of principal component analysis of Cell Painting features for 26572 control samples from 406 plates. Samples from the same plate share the same coordinate of X-axis. Note the shifts of the mean values and, to a lesser extent, differences in the ranges of values for samples from different plates.

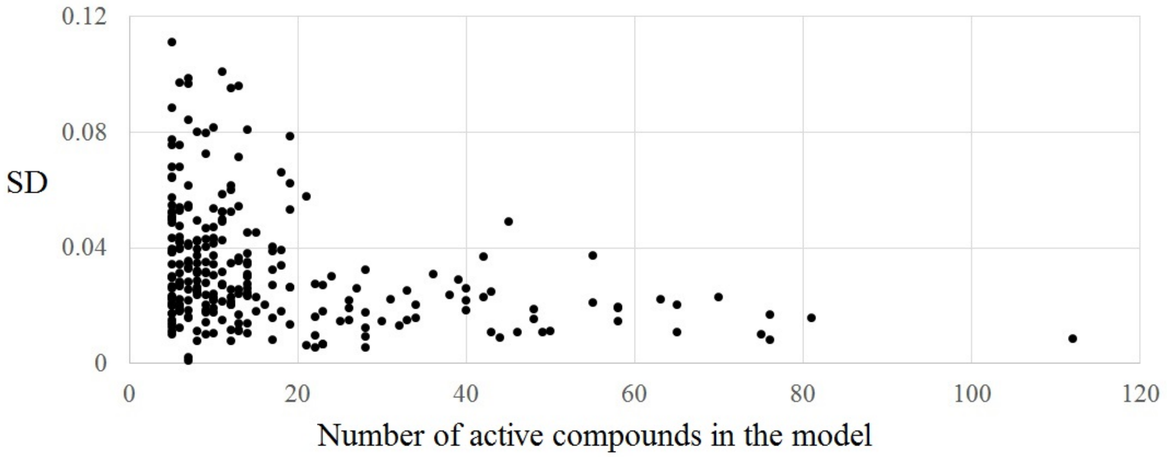


**Figure S2.** Standard deviations of AUC values of the four models built with different pre-processing and aggregation methods of Cell Painting features. Shown are results for Cell Painting data set comprising 262 MoA/Ts, where the number of active compounds per MoA/T ranges from 5 to 112.
